## Supplemental Figures S1-S8 for "The Mechanism of Replication Stalling and Recovery within Repetitive DNA"

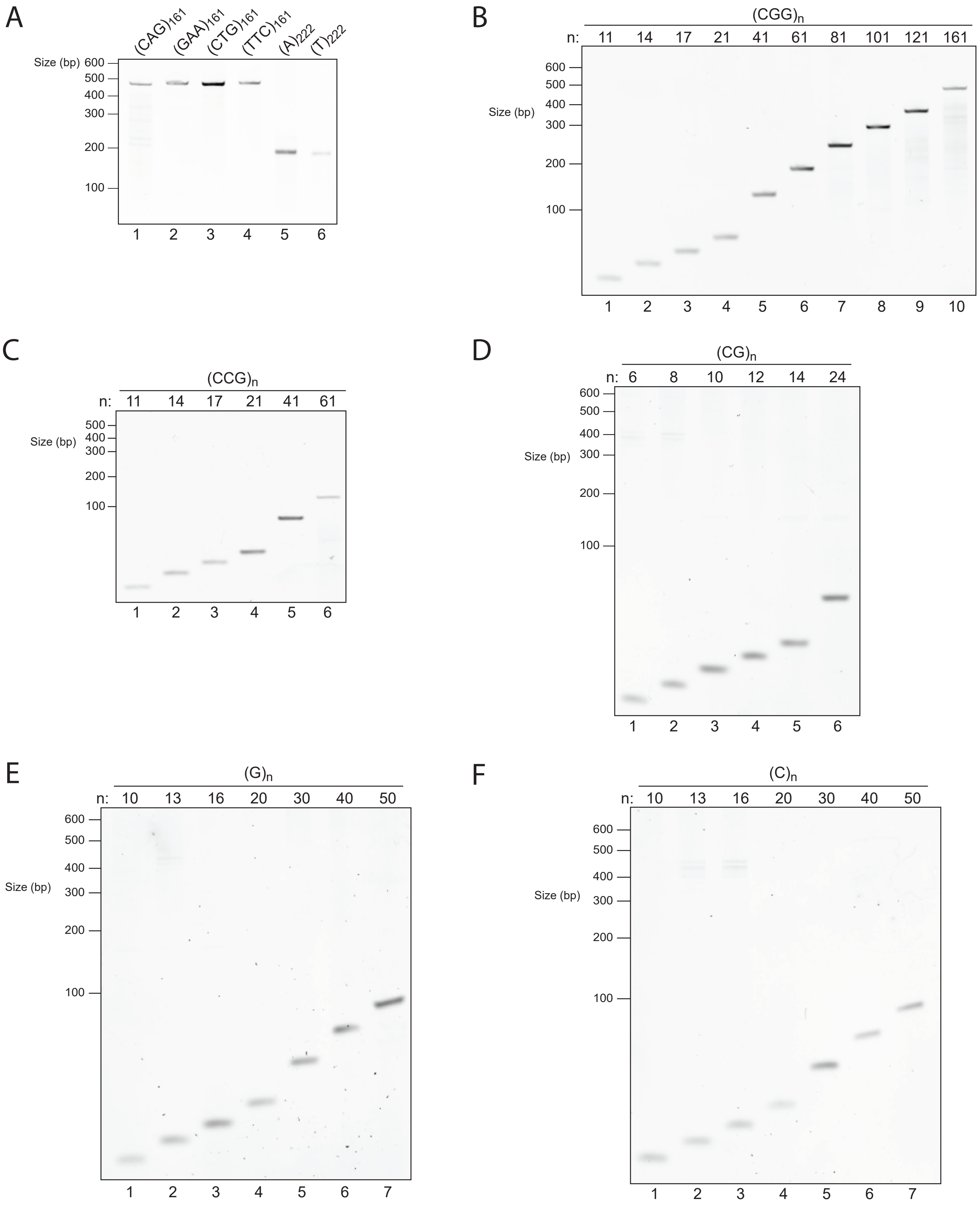


**Fig S1. Stability of repetitive sequences within final substrates used in replication assays**
To verify that repeats were stably maintained in bacteria, final preparations were cut with restriction enzymes flanking the inserts and products were analysed by TBE-PAGE and stained with SyBR gold.

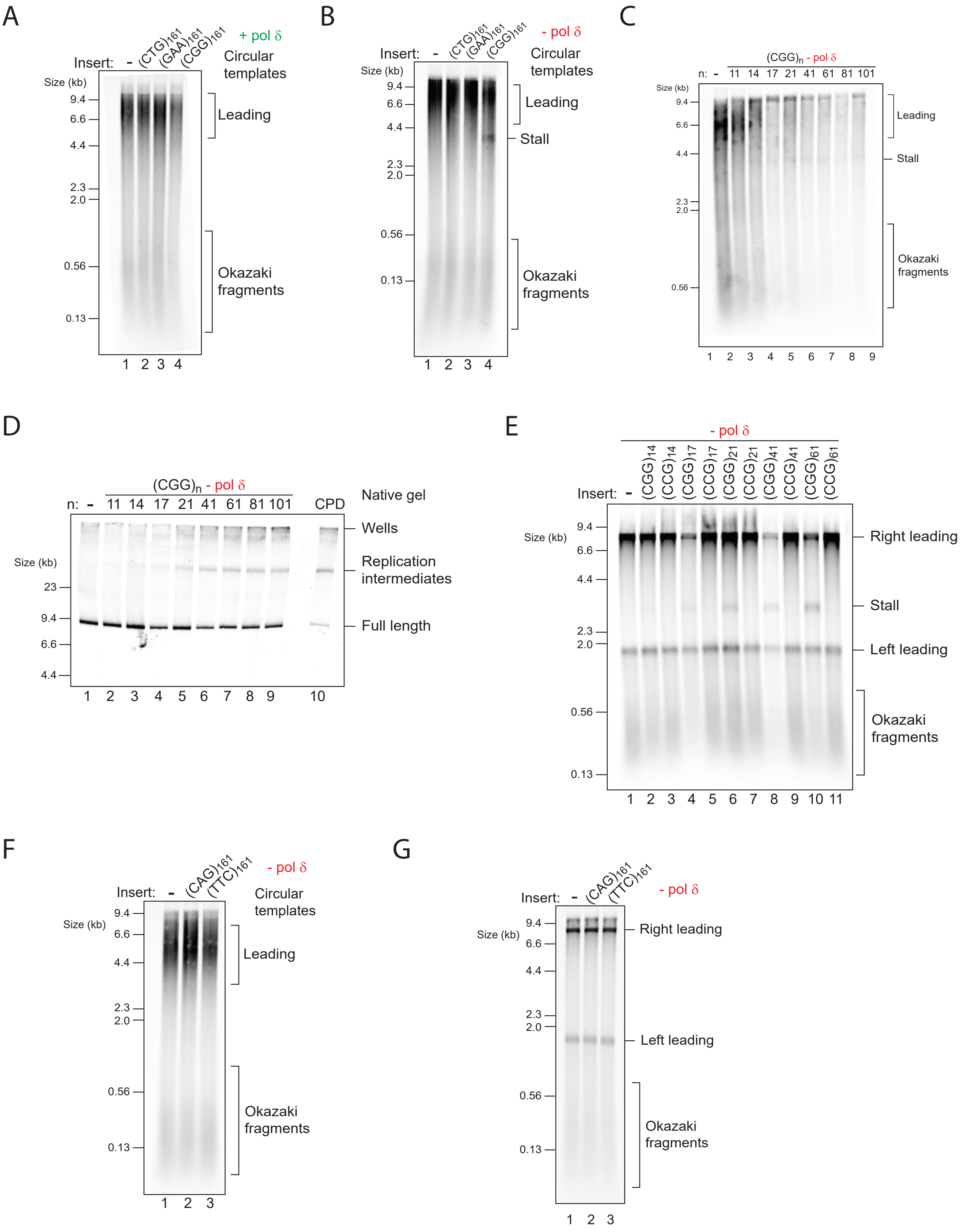


**Fig S2. Replication stalling by trinucleotide repeats**

(A and B) Replication assays with the indicated circular templates in the presence (A) or absence (B) of pol δ.
(C) Replication assays with a set of $(CGG)_n$ circular templates in the absence of pol δ.
(D) Replication assays with a set of $(CGG)_n$ linear templates in the absence of pol δ, analysed on a native gel.
(E) Replication assays comparing $(CGG)_n$ vs. $(CCG)_n$ side-by-side in the absence of pol δ.
(F and G) Replication assays with the indicated circular (F) or linear (G) substrates in the absence of pol δ.

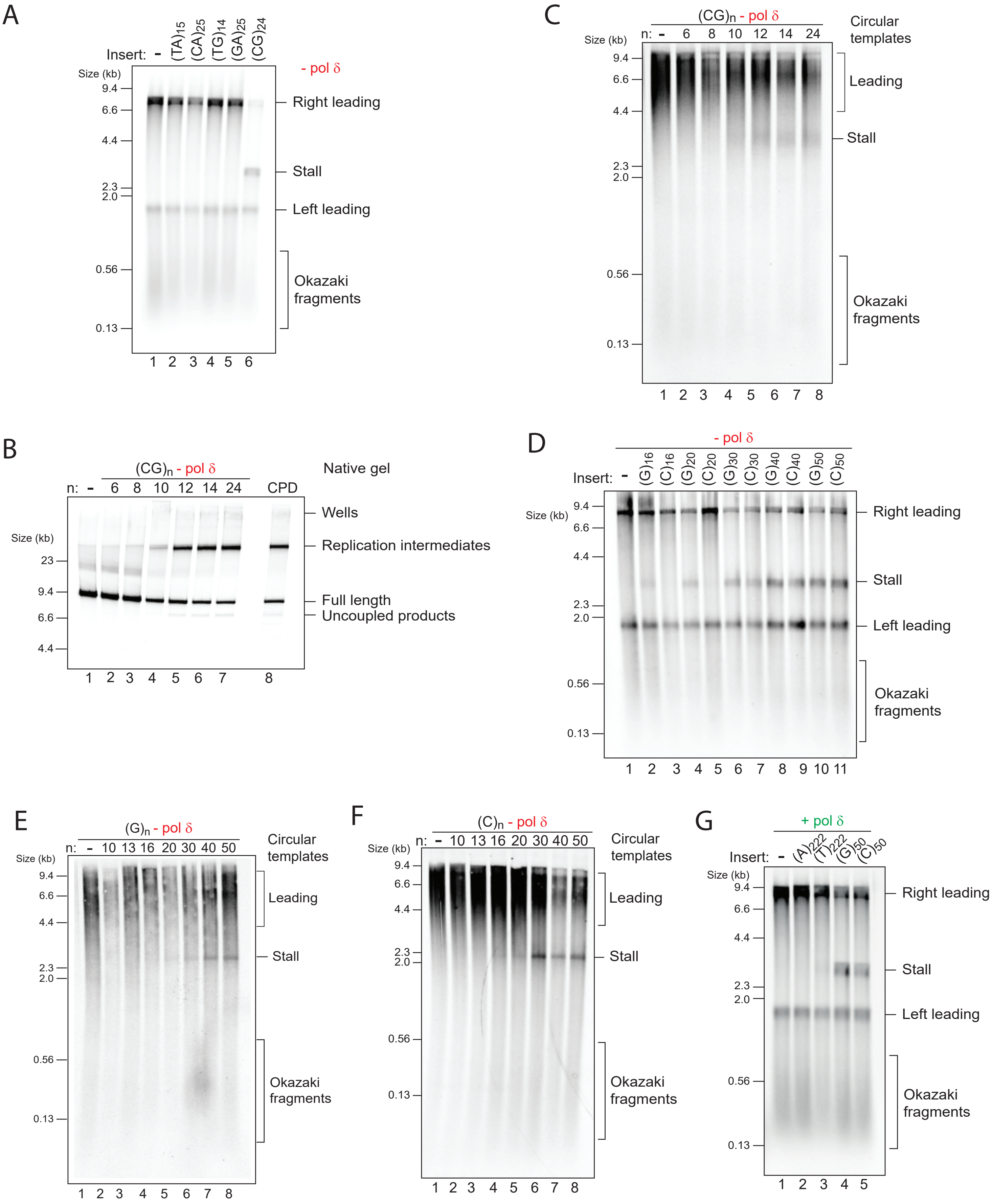


**Fig S3. Replication stalling by homopolymers and dinucleotides**
(A) Replication assays with the indicated templates in the absence of pol δ.
(B and C) Replication assays with a set of $(CG)_n$ templates in the absence of pol δ, either linear substrates analysed on a native gel (B) or circular substrates analysed on a denaturing gel (C).
(D) Replication assays comparing $(G)_n$ vs. $(C)_n$ side-by-side in the absence of pol δ.
(E and F) Replication assays with the indicated circular substrates in the absence of pol δ.
(G) Replication assays with the indicated linear substrates in the presence of pol δ.

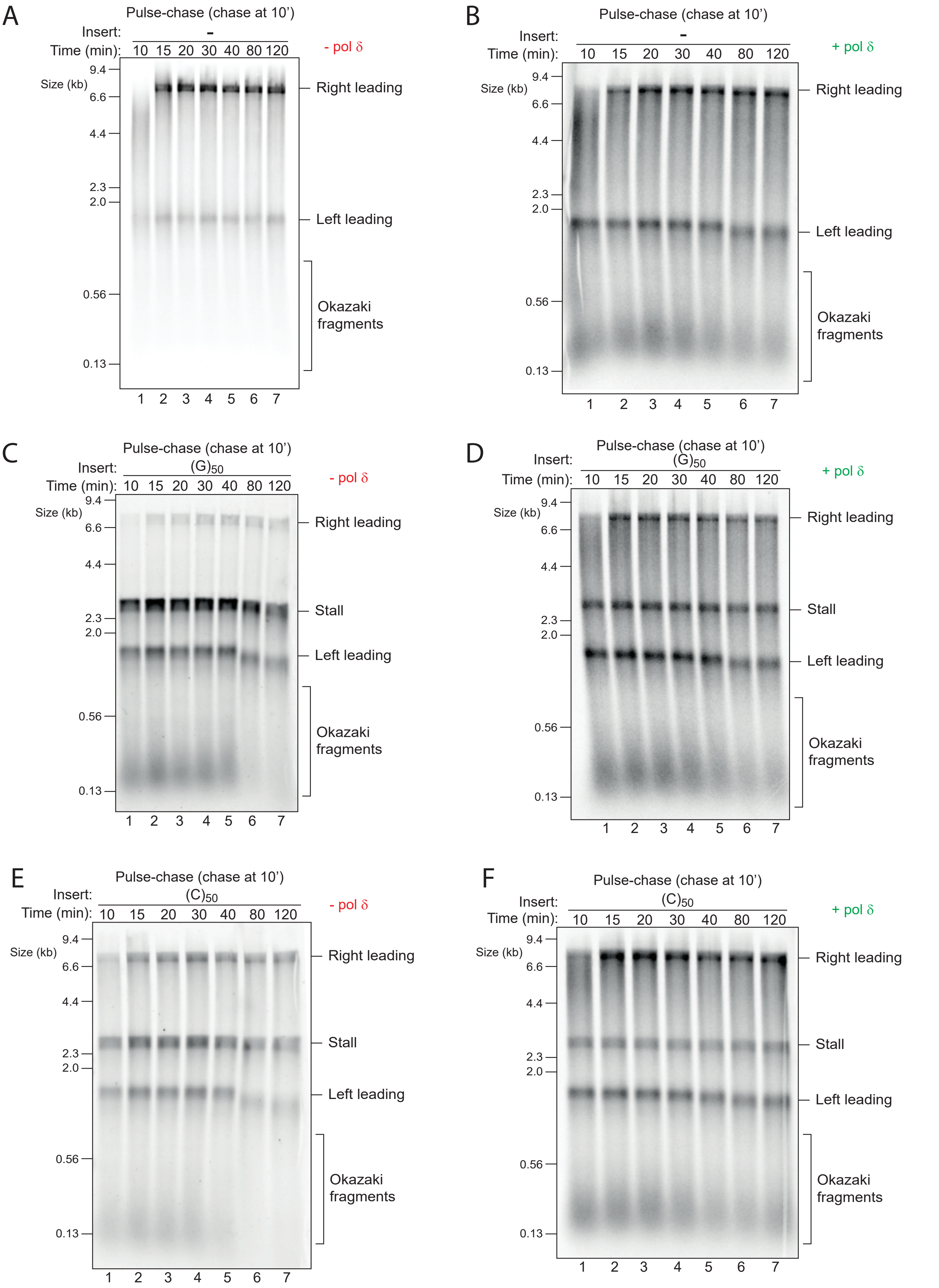


**Fig S4. Kinetics of replication stalling and recovery**

(A-F) Pulse-chase experiments with the indicated templates in the absence (A, C, E) or presence (B, D, F) of pol δ. Reactions were initiated with radiolabelled dATP for 10 min, chased with excess 'cold' dATP and samples taken at the indicated time points.

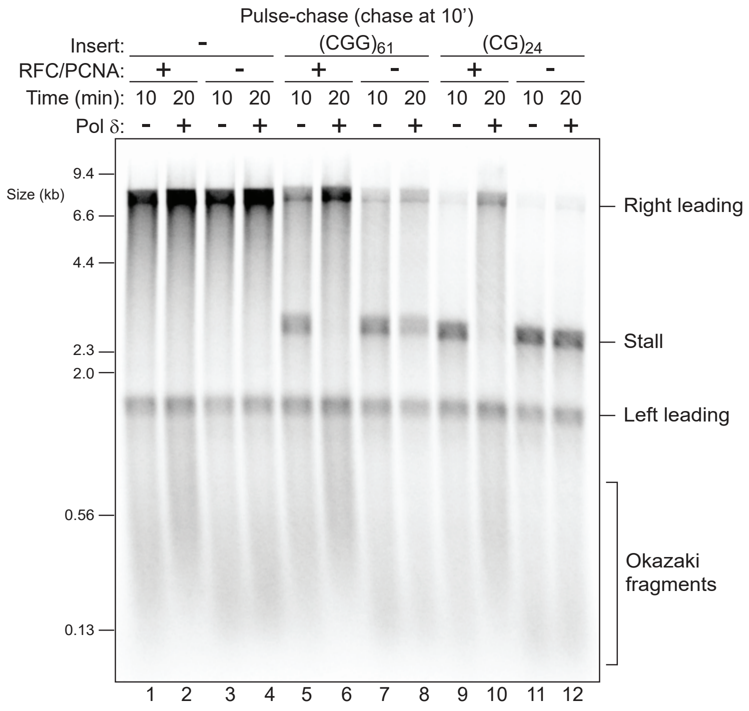


**Fig S5. RFC/PCNA are required for recovery from stalls induced by hairpin-forming sequences**
Pulse-chase experiments with the indicated templates as in Fig. 3F in the absence or presence of RFC/PCNA.

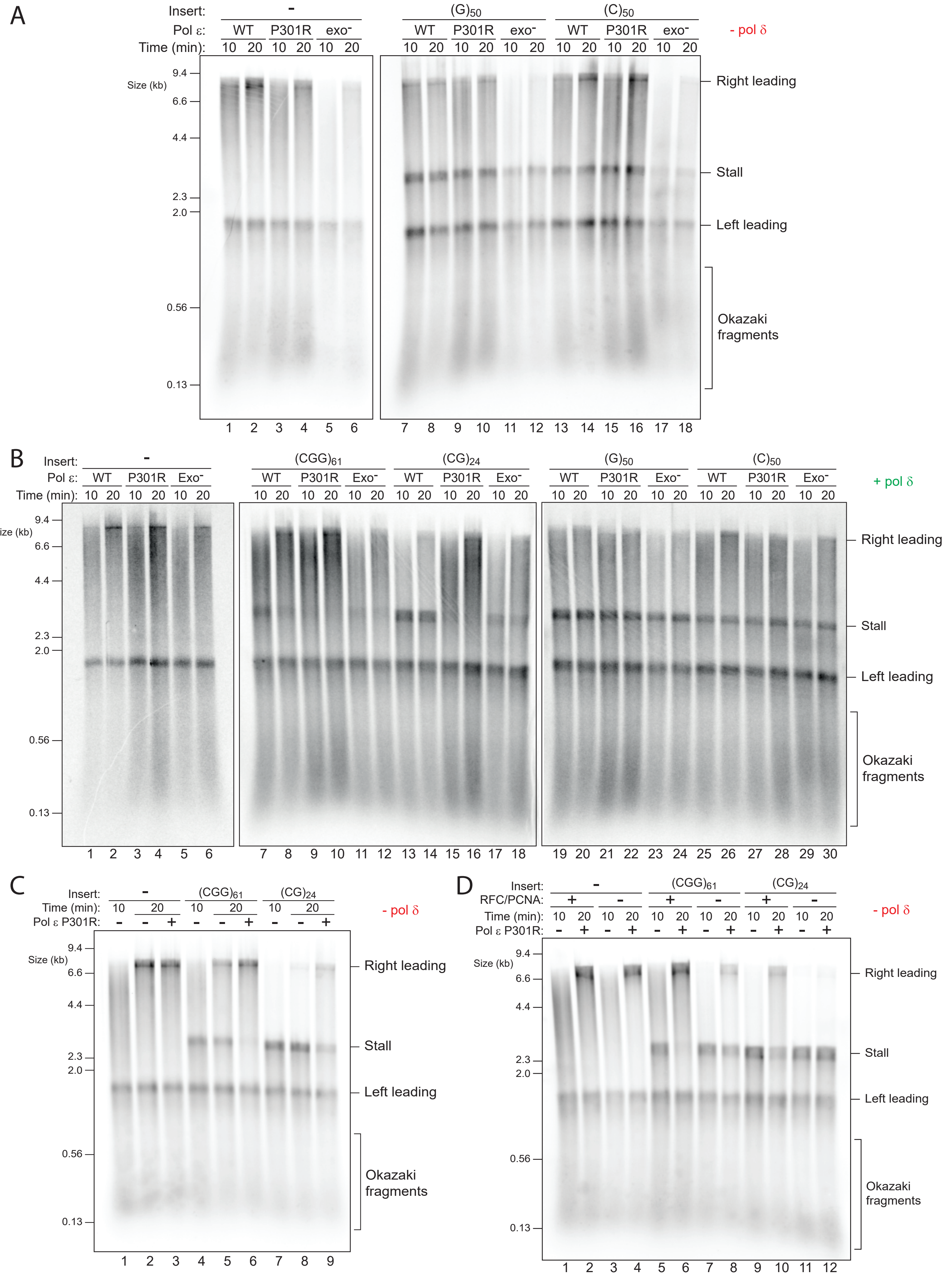


**Fig S6. Pol ε variants can drive recovery from stalls induced by hairpin-forming sequences**
(A and B) Replication assays with the indicated templates and pol ε variants in the absence (A) or presence (B) of pol δ.
(C) Pulse-chase experiment in the absence of pol δ and in the absence or presence of pol ε P301R in the chase.
(D) Pulse-chase experiment as in (C) but in the absence or presence of RFC/PCNA.

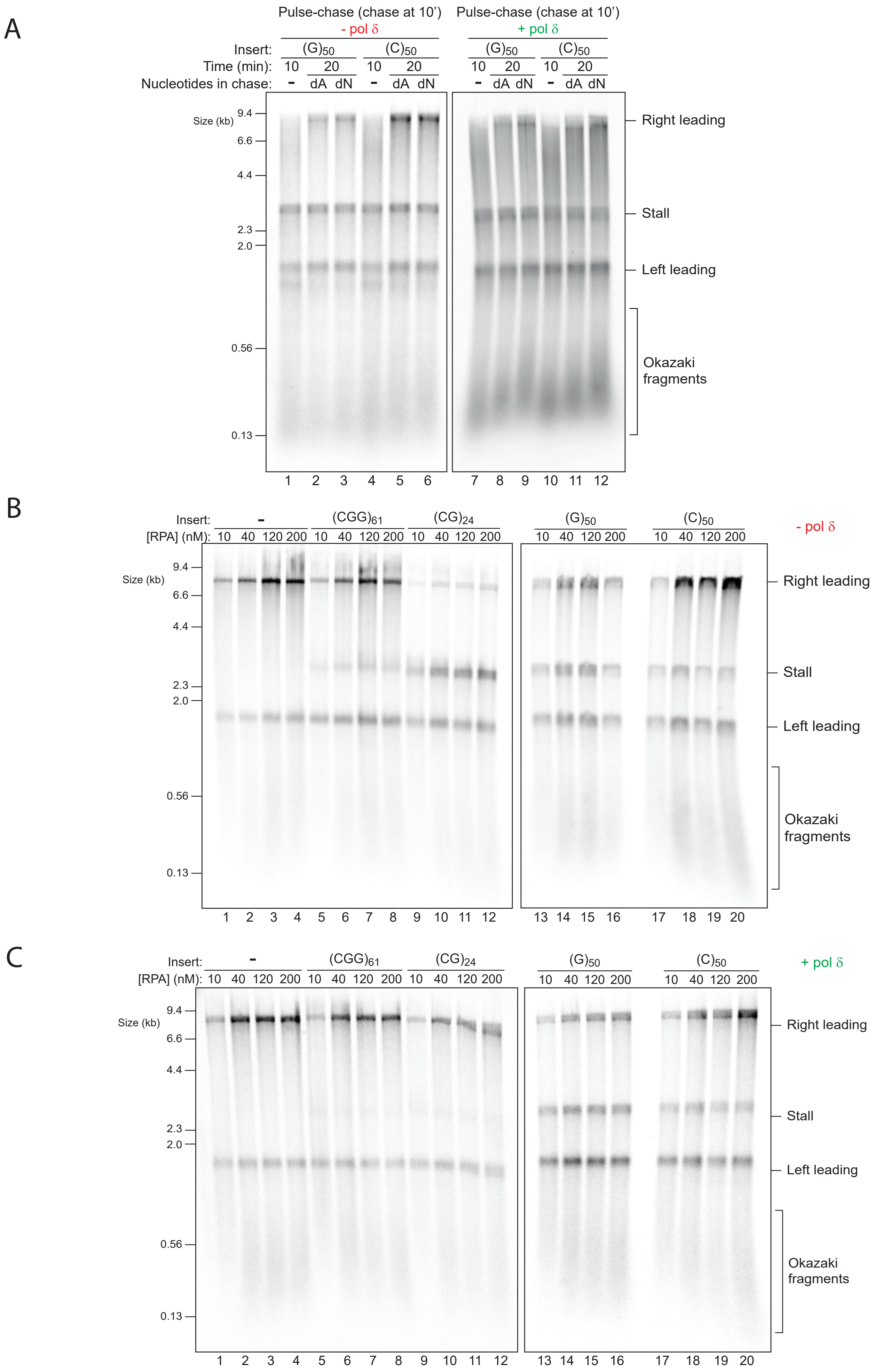


**Fig S7. Role of excess dNTPs and RPA concentration on DNA-induced stalls**

(A) Pulse-chase experiments in the absence (left panel) or presence (right panel) of pol δ. Reactions were initiated with radiolabelled dATP for 10 min and chased for another 10 min with either excess 'cold' dATP alone (dA) or with excess of all four dNTPs (dN).

(B and C) Replication assays with varying concentrations of RPA in the absence (B) or presence (C) of pol δ.

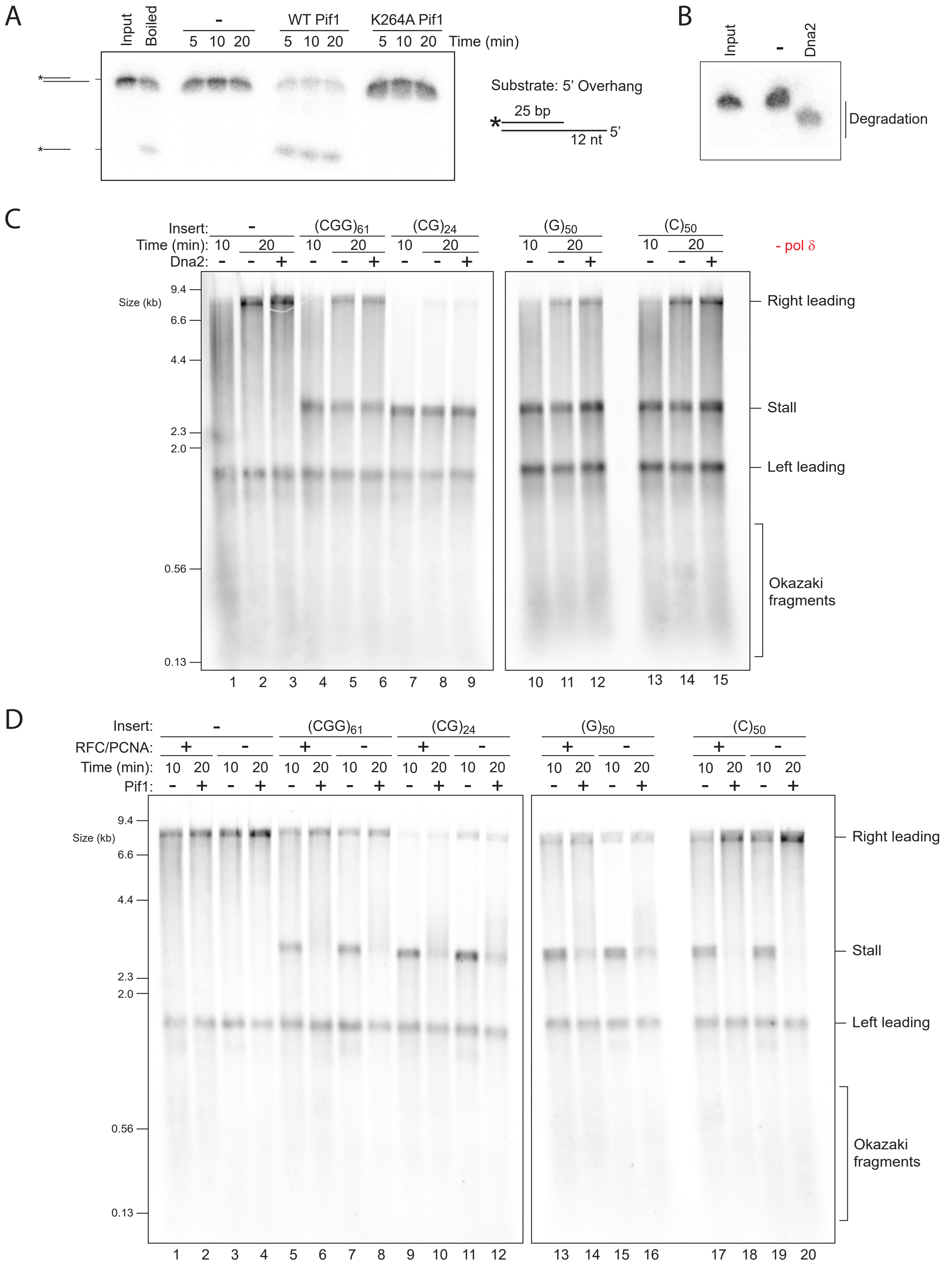


**Fig S8. WT Pif1, but not ATPase-dead Pif1 or WT Dna2, can drive efficient recovery and requires RFC/PCNA**
(A) Helicase assay shows that WT Pif1 exhibits unwinding activity whereas the K264A mutant does not.
(B) Incubation of purified Dna2 with a labelled substrate reveals robust nuclease activity.
(C) Pulse-chase experiment as in Fig. 6B but with Dna2 instead of Pif1 added to the chase.
(D) Pulse-chase experiment as in Fig. 3F but in the absence or presece of RFC/PCNA.
